# Lipid extraction from dried blood spots and dried milk spots for high throughput lipidomics

**DOI:** 10.1101/2020.06.22.165514

**Authors:** Samuel Furse, Albert Koulman

**Affiliations:** Core Metabolomics and Lipidomics Laboratory, Wellcome Trust-MRC Institute of Metabolic Science, University of Cambridge, Box 289, Cambridge Biomedical Campus, Hills Road, Cambridge CB2 0QQ, UK

**Keywords:** Lipidomics, Dried milk spots, dried blood spots, lipid, extraction efficiency

## Abstract

Dried blood spots (DBS) and dried milk spots (DMS) represent convenient matrices for collecting and storing human samples. However, the use of these sample types for researching lipid metabolism remains relatively poorly explored, and especially the efficiency of lipid extraction is unclear. A visual inspection of punched DBSs after standard extraction suggests that the samples remain largely intact. DMSs comprise a dense aggregate of milk fat globules on one side of the card, suggesting that the lipid fraction may be physically inaccessible. This led us to the hypotheses that decoagulating may facilitate lipid extraction from both DBSs and DMSs. We tested decoagulation using a mixture of strong chaeotropes (guanidine and thiourea) in both DBS and DMS in the context of high throughput lipidomics (96/384w plate). Extraction of lipids from DMSs was tested with established extractions and one novel solvent mixture in a high throughput format. We found that exposure of DBSs to chaeotropes facilitated collection of the lipid fraction but was ineffective for DMSs. The lipid fraction of DMSs was best isolated without water, using a mixture of xylene/methanol/isopropanol (1:2:4). We conclude that decoagulation is essential for efficient extraction of lipids from DBSs and that a non-aqueous procedure using a spectrum of solvents is the best procedure for extracting lipids from DMSs. These methods represent convenient steps that are compatible with the sample structure and type, and with high throughput lipidomics.

**Highlights:**

- The efficiency of lipid extractions on dried milk and dried blood spots was tested
- The number of lipid variables and the total signal strength were used as objective measures
- Decoagulation of dried blood spots improved extraction efficiency
- A mixture of xylene, methanol and isopropanol isolates the lipid fraction best from DMSs
- An aqueous extraction using dichloromethane was the most efficient method for isolating lipids from DBSs

## Introduction

Dried blood spots (DBSs) and dried milk spots (DMSs) represent convenient matrices for collecting and storing human samples. This has led to tests of their use for measuring several metabolites in a range of studies^1–7^. DBSs in particular have reached broad use in medicinal chemistry and toxicokinetic research as well as lipidomics and metabolomics studies. This sample format has also been found to be convenient for collecting samples from infants, examples including the screening of in-born metabolic disease^8–11^. More recently, measurements of endogenous lipids isolated from DBSs have enabled testing of hypotheses about human metabolism, and demand for this work is expected to increase with an increasing number of trials and other studies being planned and executed^12–15^. The increasing demand for lipidomics in metabolic studies, and the convenience of dried blood spot collection by participants at home, make DBSs and DMSs convenient sample types for larger studies focused on high throughput lipidomics.

Several studies have used DBSs for testing hypotheses about lipid metabolism^7, 16^ and work investigating the analytical limits of this sample type for lipidomics has also been reported^5, 17, 18^. Early work on DMSs has also been reported^19^. These studies have been made possible by considerable progress in laboratory infrastructure and methods for the high throughput lipidomics, including Direct Infusion Mass Spectrometry (DI-MS)^16, 20, 21^. The increasing demand for this sample type in high throughput studies, and the desire for high quality data, led us to answer unresolved questions about the analytical issues associated with using either DBSs or DMSs.

The first of these concerns the 3D structure of dried sample spots. DBSs typically form from blood soaked through the card, forming a dry and tightly coagulated mass. This differs markedly from the structure of DMSs, that present as a dried film of milk fat globules on one side of the filter paper. The coagulation typical of the structure of DBSs preserves much of the lipid fraction away from oxygen and water, but it is unclear whether the full lipid fraction of the spot can be isolated when the proteins are not decoagulated. In the case of DMSs, it is not clear whether part of the lipid fraction in DMSs also becomes inaccessible as it dries. We therefore sought to test extraction efficiency of the lipid fraction from dried milk and dried blood spots. Secondly, little work has been published on DMSs and thus the handling of this sample matrix is not fully described.

In order to improve extraction efficiency, we tested the use of a proven chaeotropic buffer for decoagulating DBSs and DMSs. Treatment with this buffer has previously been found to facilitate preparation of biological samples for lipid extraction^22^. This led to considerable improvement in the extraction efficiency of DBSs but no improvement of that from DMSs, leading us to suppose that lipids were not inaccessible in DMSs. We therefore tested several solvent systems for extracting lipids from DMSs in order to identify a method that was convenient for the simultaneous extraction of large numbers of small samples and for profiling as many of the lipid components present as possible.

To assess the extractions objectively we used Direct Infusion Mass Spectrometry (DIMS)^20, 21^ to quantify both the number of lipid isoforms (variables) identified and the total signal strength of lipid variables. This answers concerns raised about how lipid extraction procedures are compared, improving on speculative tests that are inconclusive for analytical purposes ^23^. We argue that it is important to investigate and improve these methods in order to be able to choose the most appropriate method(s) for testing hypotheses about lipid metabolism. The present work can be used to facilitate the analysis of sets of hundreds or thousands of DBSs and/or DMSs for metabolic studies.

## Materials and Methods

### Reagents and standards

Solvents were purchased from *Sigma-Aldrich Ltd* (Gillingham, Dorset, UK) of at least HPLC grade and were not purified further. Lipid standards were purchased from Avanti Polar lipids (Alabaster, AL; through Instruchemie, Delfzijl, NL) and used without purification. Consumables and pooled synthetic blood products were purchased from Sarstedt AG & Co (Leicester, UK), Thermo Fisher (Hemel Hempstead, Herfordshire, UK). Commercially available pooled human serum, synthetic and anonymised (unidentifiable) human blood were used to prepare dried blood spots (DBS).

### Sample processing

Blood or pasteurised animal milk/almond drink was spotted onto commercially available Guthrie spot cards and left to dry (2-16 h) before use. Cards were stored in air-tight bags with minimum gaseous volume at −80°C and in the presence of a mild drying agent. Cards were punched with a hand-held puncher (3.2mm disc) into the extraction plate (glass-coated 2.4 mL/well 96w plate; Plate+™, Esslab, Hadleigh, UK). DBSs were treated with a prepared solution (60 μL/well) of guanidinium chloride (6 M) and thiourea (1.5 M) in the extraction plate (plastic sealing tape) at ~10 °C for the appropriate period.

### Quality control

QC samples were used to establish which variables’ signal strength correlated with their concentration. Three QC levels were used, 0.25, 0.5 and 1.0× of the total (20 μL). The reference material used was designed to be as broad as possible. It comprised equal parts of an homogenate of mouse placenta^22^, pooled human blood serum and an infant-formula/animal milk preparation. The latter mixture comprised infant formula based on caprine, bovine and soya material, made up to a solution (15 mg mL^−1^) using Jersey milk described previously^24, 25^. The three solutions were mixed and stored in aliquots until use.

### Isolation of lipid fractions

#### DMT

This procedure was similar to a high throughput technique described recently^21, 22^. Punched DBS/DMSs (3.2mm) were placed along with blank and QC samples in the wells of a glass-coated 2.4 mL/well 96w plate (Plate+™, Esslab, Hadleigh, UK). Methanol (150 μL, HPLC grade, spiked with Internal Standards, See *Table S1*), was added to each of the wells, followed by water (500 μL) and a mixture of solvents (500 μL) comprising dichloromethane and methanol (3:1) doped with triethylammonium chloride (500 mg L^−1^). The mixture was agitated (96 channel pipette) before being centrifuged (3·2k × *g*, 2 min). A portion of the organic solution (20 μL) was transferred to a high throughput plate (384w, glass-coated, Esslab Plate+™) before being dried (N_2 (g)_). The dried films were re-dissolved (TBME, 30 μL/well) and diluted with a stock mixture of alcohols and ammonium acetate (90 μL/well; propan-2-ol : methanol, 2:1; CH_3_COO.NH_4_ 7·5 mM). The analytical plate was heat-sealed and run immediately.

#### TBME

This procedure was similar as possible to the original protocol for extracting lipids from biological samples^26^. Punched DMSs (3·2 mm) were placed along with blank and QC samples in the wells of a glass-coated 2.4 mL/well 96w plate (Plate+™, Esslab, Hadleigh, UK). Methanol (150 μL, HPLC grade, spiked with Internal Standards, See *Table S1*), was added to each of the wells, followed by water (500 μL) and TBME (500 μL). The mixture was centrifuged (3·2k × *g*, 2 min). A portion of the organic solution (20 μL) was transferred to a high throughput plate (384w, glass-coated, Esslab Plate+™) before being dried (N_2 (g)_). The dried films were re-dissolved (TBME, 30 μL/well) and diluted with a stock mixture of alcohols and ammonium acetate (90 μL/well; propan-2-ol : methanol, 2:1; CH_3_COO.NH_4_7·5 mM). The analytical plate was heat-sealed and run immediately.

#### XMI

This procedure has not been described before. Punched DMSs (3·2 mm) were placed along with blank and QC samples in the wells of a glass-coated 2.4 mL/well 96w plate (Plate+™, Esslab, Hadleigh, UK). A prepared mixture of solvents, xylene/methanol/isopropanol (1:2:4, 500 μL, methanol spiked with Internal Standards, See *Table S1*), was added to each of the wells. The mixture was agitated (gentle orbital mixer, 5 min) before a portion of the organic solution (20 μL) was transferred to a high throughput plate (384w, glass-coated, Esslab Plate+™). This was then dried before being re-dissolved (TBME, 30 μL/well) and diluted with a stock mixture of alcohols and ammonium acetate (90 μL/well; propan-2-ol : methanol, 2:1; CH_3_COO.NH_4_ 7·5 mM). The analytical plate was heat-sealed and run immediately.

### Mass Spectrometry

#### Instrument

Samples were infused into an Exactive Orbitrap (Thermo, Hemel Hampstead, UK), using a Triversa Nanomate (Advion, Ithaca US). Samples were ionised at 1·2 kV in the positive ion mode. The Exactive started acquiring data 20 s after sample aspiration began. After 72 s of acquisition in positive mode the Nanomate and the Exactive switched over to negative mode, decreasing the voltage to −1·5 kV. The spray was maintained for another 66 s, after which the analysis was stopped and the tip discarded, before the analysis of the next sample. The sample plate was kept at 15 °C throughout the acquisition. Samples were run in row order.

#### Data processing

Raw high-resolution mass-spectrometry data were processed using XCMS (www.bioconductor.org) and Peakpicker v 2.0 (an in-house R script ^20^). Lists of known species (by *m/z*) were used for both positive ion and negative ion mode (~8·5k species). Variables whose mass deviated by more than 9 ppm from the expected value, had a signal/noise ratio of <3 and had signals for fewer than 75% of samples were discarded. The correlation of signal intensity to concentration of human placenta, mouse liver, human serum and pooled human seminal plasma samples as QCs (0.25, 0.5, 1.0×) were used to identify the lipid signals the strength of which was linearly proportional to abundance (threshold for acceptance was a correlation of 0.75). Signals were compared using the raw signal strength. Normalisation of signal strength was used for assessing the coefficient of variance for individual lipid variables. The signal for an individual variable was divided by the sum of signals for that sample and expressed per mille (‰). Zero values were interpreted as not measured.

### Statistical analyses

The analysis was structured according to a prepared analysis plan. Uni- and bivariate analyses were carried out using Excel 2016. Multivariate analyses were run using MetaboAnalyst 4.0 ^27^. Abundance of lipid(s) is shown as mean ± standard deviation of relative abundance unless otherwise stated. Relative abundance was calculated based on the proportion of an individual signal as a fraction of the whole.

## Results

### Dried Blood Spots (DBS)

We tested the hypothesis that greater efficiency of lipid extraction from was possible with a decoagulated DBS using a powerful mixture of chaeotropes, guanidine and thiourea (GCTU, see *Methods*). Visual assessment of DBSs exposed to GCTU for 16h indicated that the spot was at least partly dismantled. However, visual inspections alone were not enough to establish the optimum exposure to the buffer for the best analytical results. We therefore exposed 3.2 mm discs of DBSs to GCTU for up to 48h and measured both how many variables were isolated and also at what point the signal strength was highest. This was done in both positive ionisation mode (mainly triglycerides and zwitterionic phospholipids) and negative ionisation mode (mainly zwitterionic and anionic phospholipids). This time course is shown in Fig. 1.

**Fig. 1.**
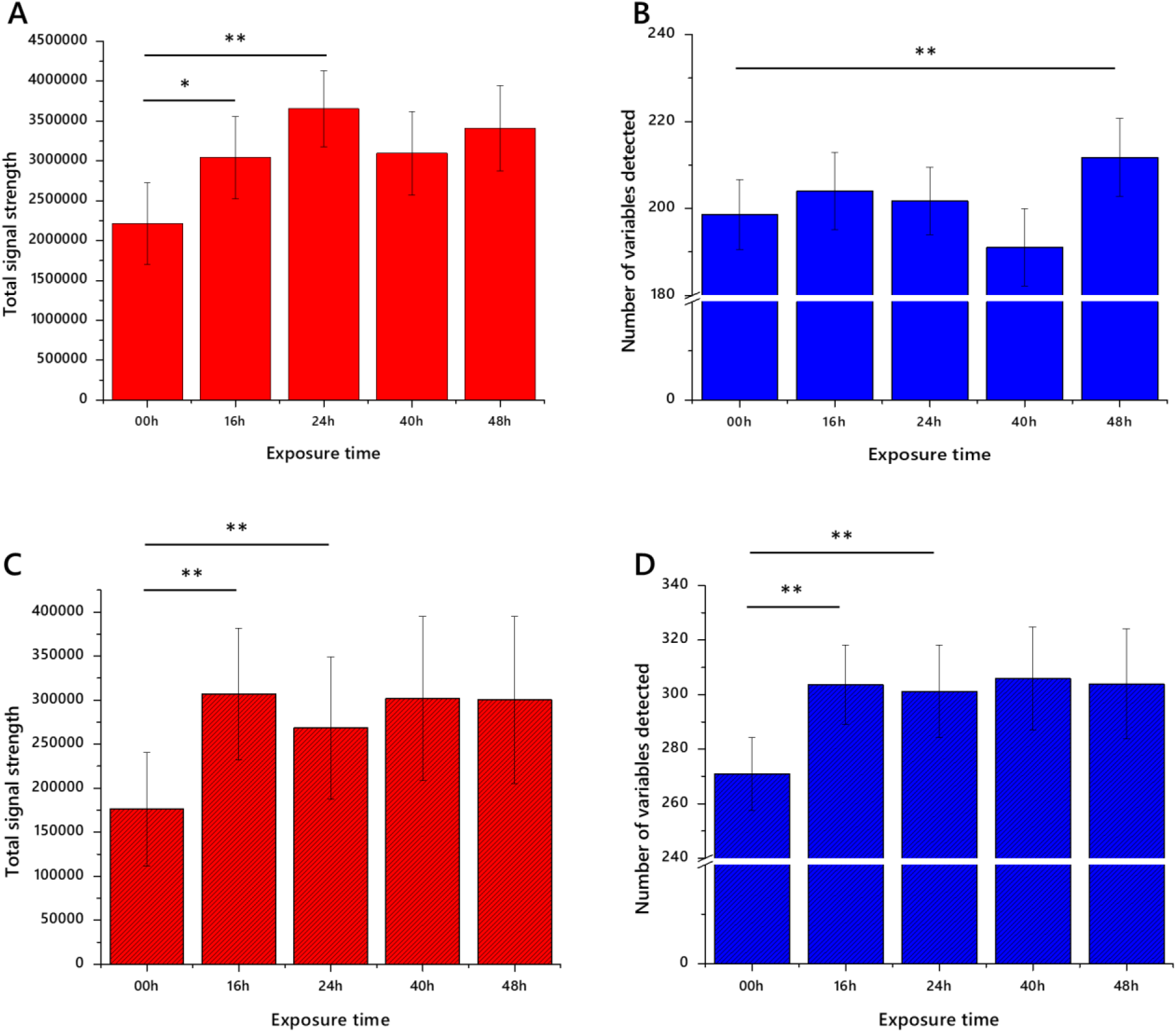
Signal strength and number of variables identified in lipid extracts of dried blood spots exposed to a guanidine-thiourea buffer. Panel A, total signal strength of QCed variables; B, number of QCed variables. Error bars show standard deviation. *n* = 16 measurements per group. \**p* <0.01, \*\**p* <0.001. The control (00h) time point represents the present method used (no chaeotropes, exposure to water shortly before exposure to organic solvents).

This experiment, using blood spots generated from commercially available human erythrocytes and plasma, showed that total signal strength in positive mode increased in the first 24h but not thereafter (Fig. 1A). There was a slight (5%) increase in the number of variables between 24 and 48h but this was not statistically significantly increased (Fig. 1B). This is reflected in the results from the negative ionisation mode, however there is little change in either signal strength (Fig. 1C) or number of variables (Fig. 1D) after 16h. These data suggested to us that 24h was the optimum time to minimise the time needed for dismantling the spots with maximum extraction efficiency.

However, we note that a longer extraction procedure might have introduced noise. This led us to characterise the precision of the modified method. The precision of this method was tested visually (a principal component analysis, PCA, Fig. 2) and quantitatively (coefficient of variance, CV, Fig. 3). In order to gain a clear insight into the performance of this method, DBSs from three healthy volunteers (cards 1-3) and the synthetic blood (card 4) used above were used.

**Fig. 2.**
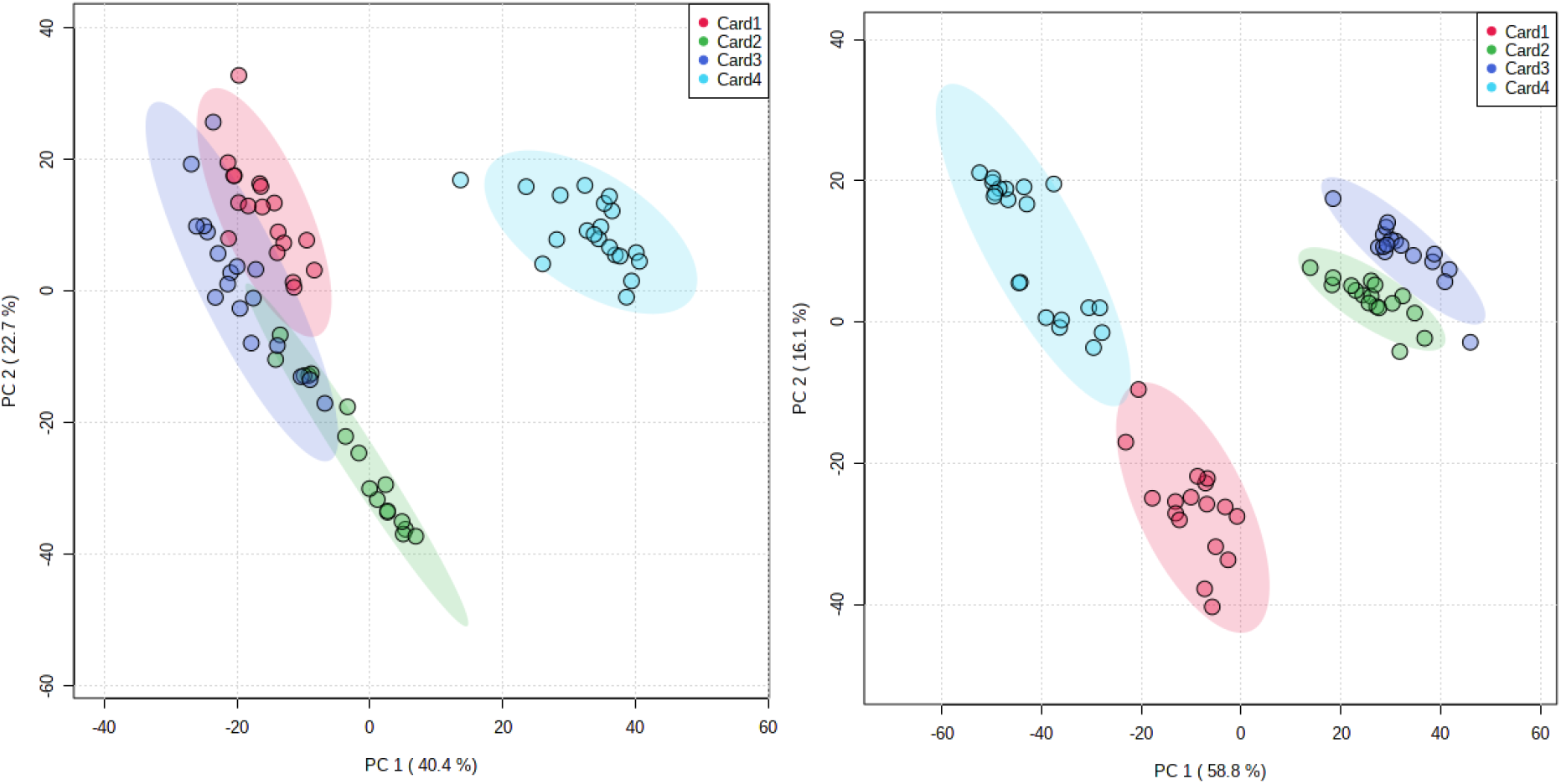
Principal component analysis of dried blood spots prepared from fresh blood from three healthy volunteers (Cards 1-3) and synthetic human blood (Card4). Panel A, positive ionisation mode (*n* = 203 variables); B, negative ionisation mode (*n* = 316 variables). Only variables that have 75% or more non-zero values were used. *n* = 20 measurements per group.

**Fig. 3.**
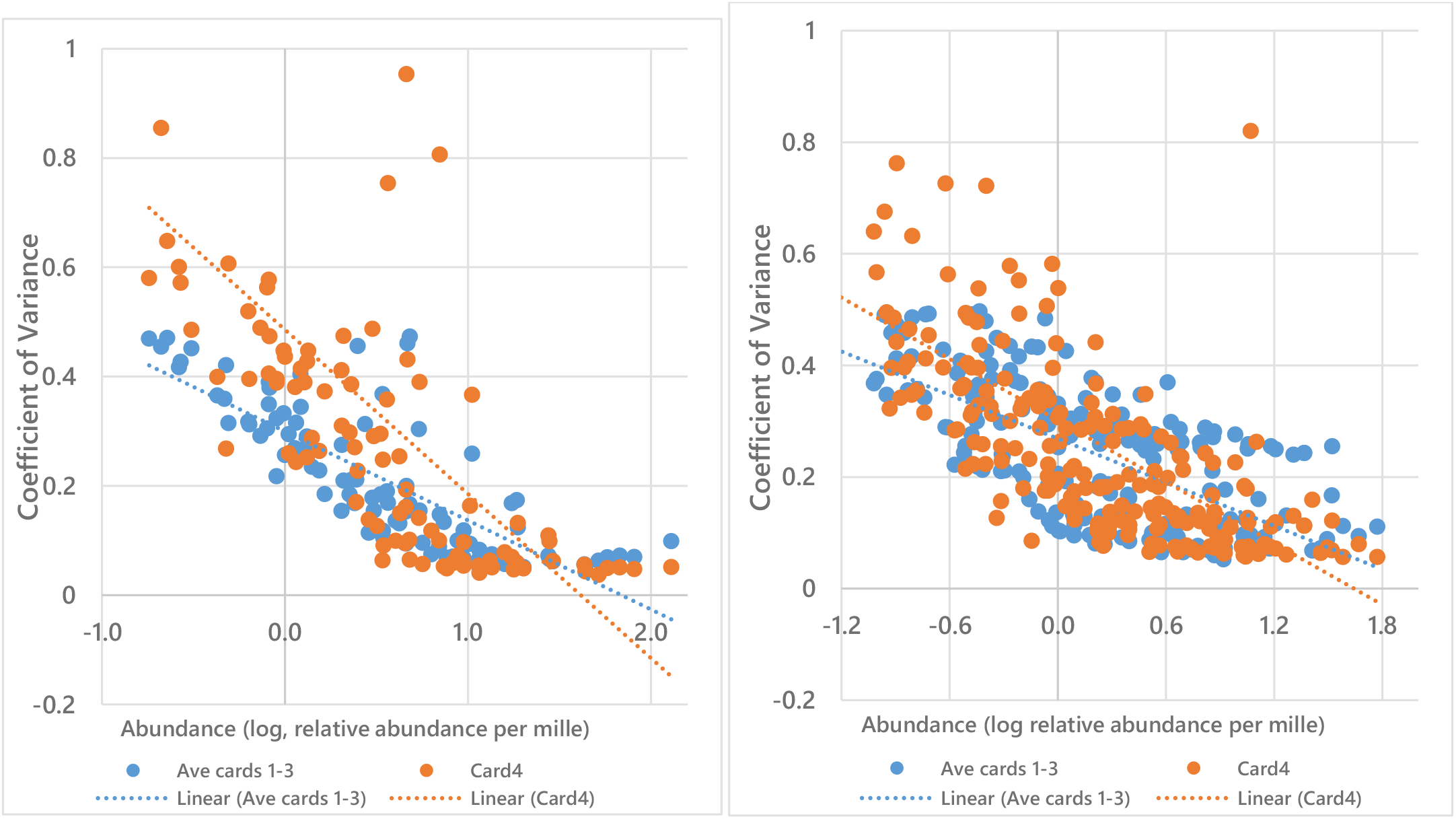
Relationship between CV and lipid abundance. Panel A, +ve mode (*n* = 99 variables); B, +ve mode (*n* = 212 variables). Variables with a CV <50% are plotted. Cards1-3 were prepared from blood from healthy volunteers. Card4 was prepared from commercially available, synthetic human blood.

A PCA showed that the four samples separated and grouped easily, particularly clearly in negative ionisation mode (Fig. 2). The coefficient of variance of the total signal was 24% (+/−4%) in negative ionisation mode and 23% (+/−1%) in positive ionisation mode. Around two thirds of variables had a CV below 50% (Table 1). Plotting the relationship between coefficient of variance and relative abundance of variables (Fig. 3) showed that the lowest abundance variables were characterised as having the highest CVs, and thus this method was consistent in terms of the most important lipid variables. It also showed that the synthetic blood used differed analytically from that of from that of healthy volunteers. The improved signal strength and increase in number of lipid variables observed, the relatively small variation in repeated measurements and the low CV suggest that treating DBSs with GCTU in advance of extraction of the lipid fraction is beneficial for high throughput lipidomics.

**Table 1.**
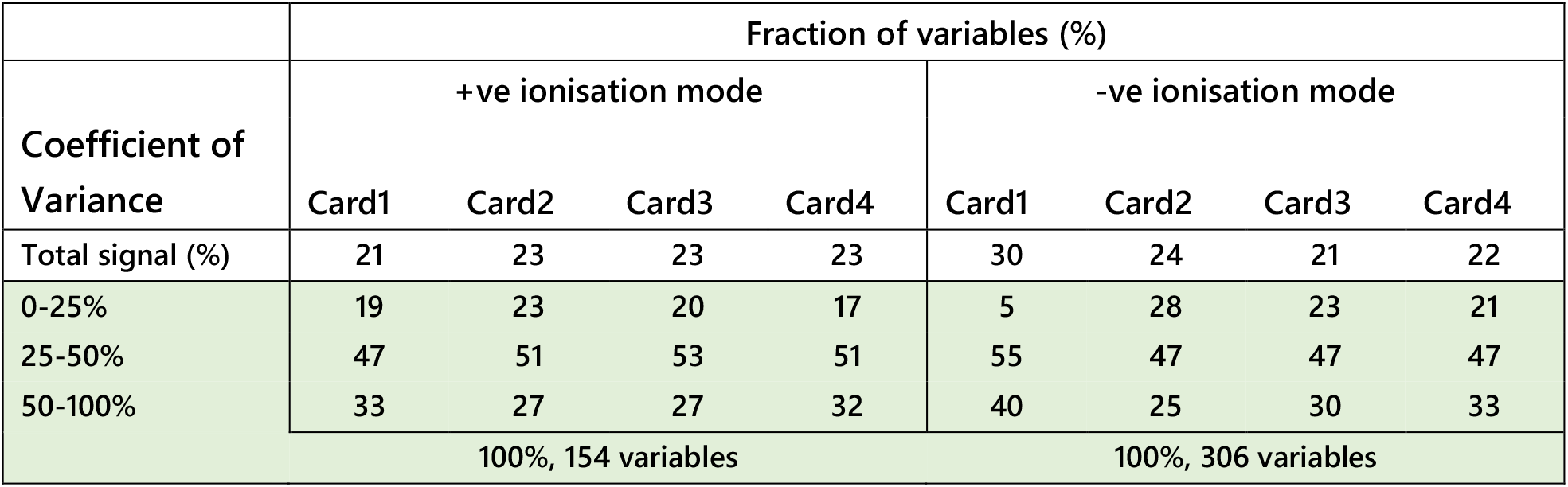
Coefficient of variance of lipid variables from dried blood spots. Dried blood spots were prepared from fresh blood donated from three healthy volunteers (Cards 1-3) and synthetic human blood (Card4). Only variables with fewer than 15% 0 values were used, made possible by the relatively homogenous nature of the . *n* = 20 measurements per group.

### Dried Milk Spots (DMSs)

DMSs are an equally convenient sample matrix to that of DBSs, but are less commonly applied or well understood. As the structure of DBSs makes access to the lipid fraction difficult, we tested whether decoagulants improved signal strength or the number of variables observed for DMSs as well. As a visual appraisal of DMSs suggested that milk fat globules may aggregate on one face of the card, we also tested different solvent mixtures in order to determine which optimised isolation of the lipid fraction from this dense aggregation. We tested a novel extraction procedure against two methods reported for high throughput lipidomics, ‘DMT’, (dichloromethane/methanol/triethylammonium chloride, 3:1:0.005)^21, 22, 25^ and ‘TBME’ (*tertiary*-butylmethyl ether)^26^, both of which used an aqueous wash as part of the extraction. The novel procedure was developed based on an established method that employed a mixture of chloroform, methanol and isopropanol (1:2:4)^28, 29^. However, this was difficult to incorporate in high throughput approach due to solvent evaporation. We therefore replaced chloroform with xylene. The latter solvent is much less volatile than chloroform, has a very small dipole moment and is well-established for removing lipophilic material during the preparation of histology samples. We therefore tested ‘XMI’ (xylene/methanol/isopropanol, 1:2:4, spiked with internal standards) as a solvent mixture for extracting lipids from DMSs.

The first stage of testing lipid extraction methods on DMSs was to explore whether decoagulants (chaeotropes) were required to gain better access to the lipid fraction. We therefore tested DMT and DMT with GCTU. This test showed that despite the facility with DBSs, decoagulants do not increase either the signal strength of lipid extracts (Fig. 4A) or the number of variables identified (Fig. 4B) in the extraction of lipids from DMSs. We also wanted to test whether chlorinated, alcoholic or ethereal solvents were superior to one another for extracting from DMSs. DMT and TBME were therefore tested against one another and XMI (Fig. 4). These also showed that DMT and XMI were both superior to TBME in terms of signal strength. However, it was not clear to us whether this could be ascribed to the format of the extraction (dry/wet), and so we tested whether DMT without an aqueous wash compared favourably with XMI. It does not. Notably however, despite up to half an order of magnitude difference in the total signal strength there was no significant difference in the number of variables identified.

**Fig. 4.**
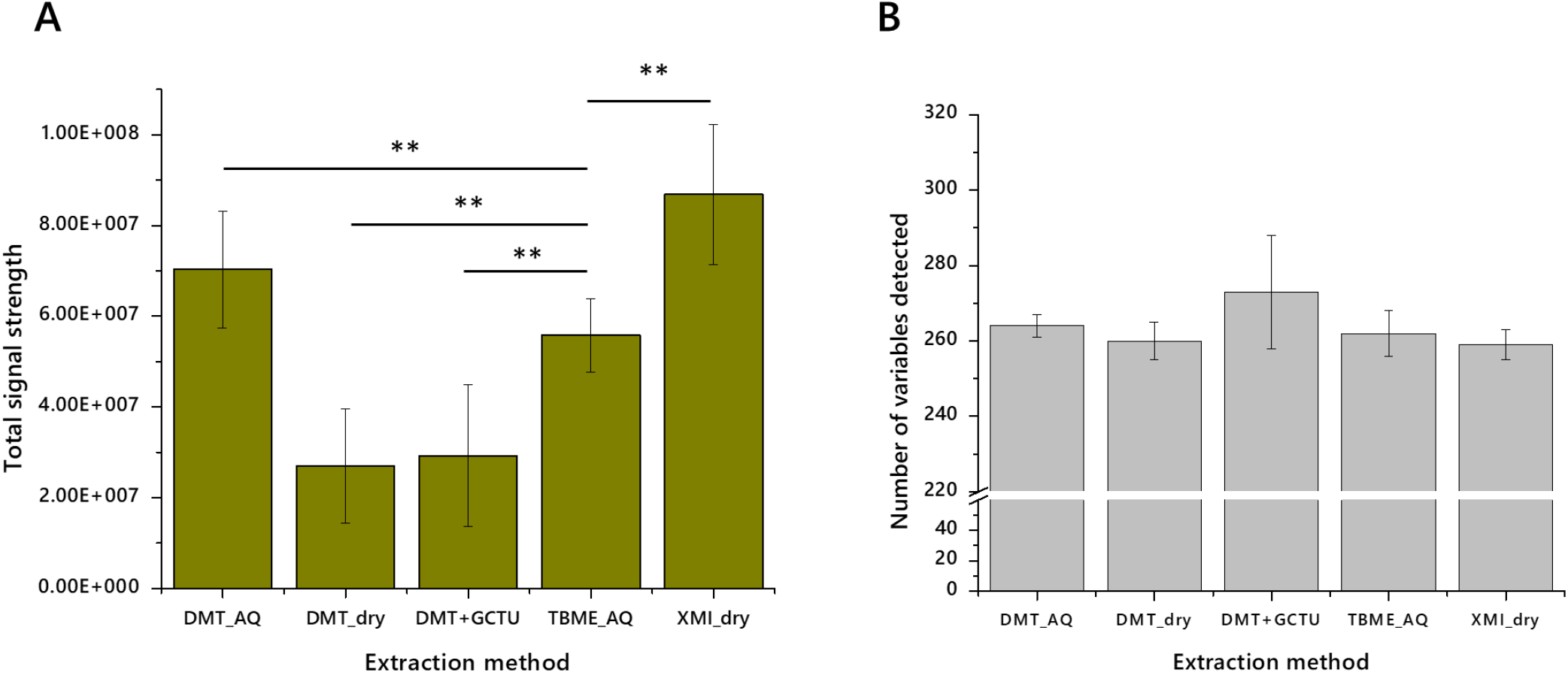
Total signal strength and the number of variables observed for a variety of lipid extraction protocols tested on dried milk spots (whole bovine milk, 3·5% fat) in positive ionisation mode. Panel A, Total signal strength based on one punched DMS; B, number of variables with non-zero signal strength. Error bars represent standard deviation. *n* = 10 measurements per group. \**p* <0.01, \*\**p* <0.001.

This led us to test the best three methods in more depth. DMT, TBME and XMI methods were used to extract lipids from a greater number (*n* = 16) samples on DMSs made from unhomogenised Jersey milk. The resulting extracts were also profiled in both positive ionisation mode (Figs 5A, B) and negative ionisation mode (Fig. 5C, D). These experiments showed once again that DMT extracted lipid fractions have a higher signal strength (implying more material, Fig. 5A, C) and also more variables in positive mode (Fig. 5B). We also found that XMI isolates more variables still further in positive ionisation mode (Fig. 5B), though without additional signal strength (Fig. 5A). The signal strength is similar for both DMT and XMI in negative ionisation mode, around twice that of TBME. This is ascribed to the greater ability of chlorinated and alcoholic solvents in to dissolve phospholipids.

**Fig. 5.**
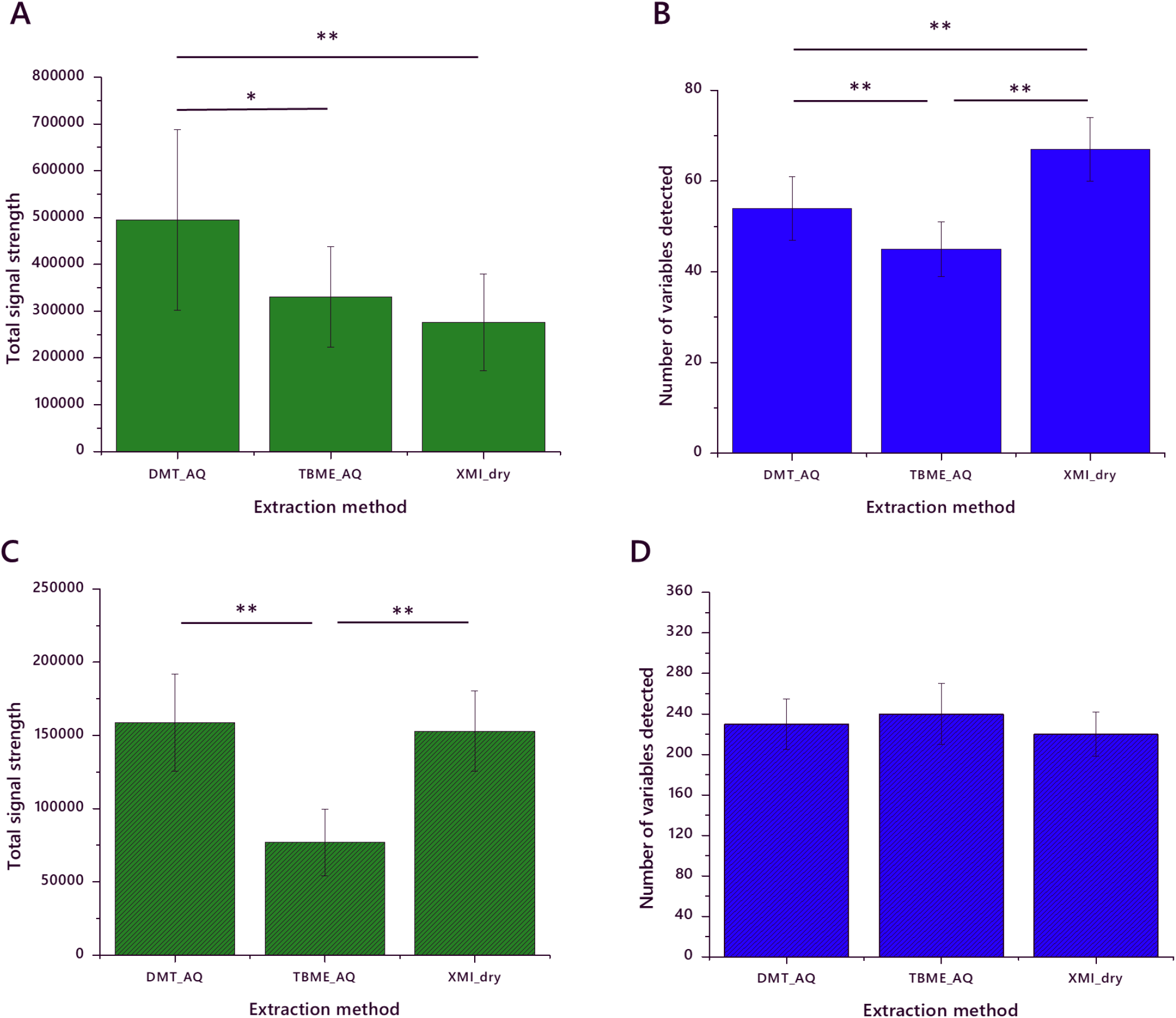
Signal strength and number of variables identified in lipid extracts of dried milk spots made from Jersey milk (5% fat). Panel A, total signal strength of lipid variables in positive mode; B, number of lipid variables in positive mode; C, total signal strength of lipid variables in negative mode; D, number of lipid variables in negative mode. Error bars show standard deviation. *n* = 16 measurements per group. \**p* <0.01, \*\**p* <0.001. DMT_AQ, extraction using dichloromethane/methanol/triethylammonium chloride (3:1:0.005) with an aqueous wash; TBME_AQ, extraction using *tertiary*-butylmethyl ether with an aqueous wash; XMI_dry, extraction using xylene/methanol/isopropanol (1:2:4) with no aqueous wash.

**Fig. 6.**
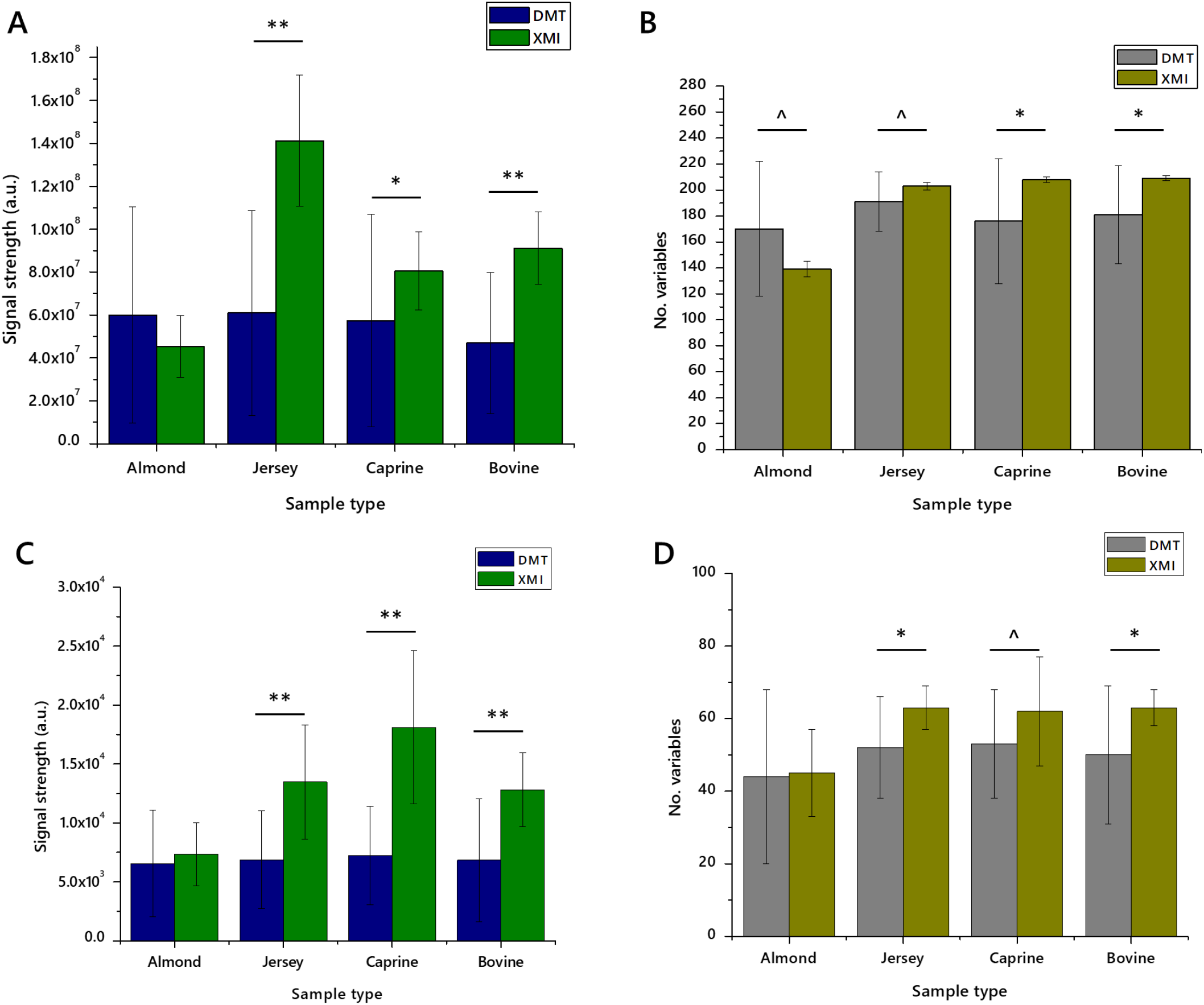
Signal strength and number of variables identified in lipid extracts of dried milk spots made from whole bovine, unhomogenised Jersey, and whole caprine milks and almond drink. Panel A, total signal strength of lipid variables in positive mode; B, number of lipid variables in positive mode; C, total signal strength of lipid variables in negative mode; D, number of lipid variables in negative mode. Error bars show standard deviation. *n* = 20 measurements per group. ^ *p* <0.05, \**p* <0.01, \*\**p* <0.001. DMT, extracted with a mixture of dichloromethane, methanol and triethylammonium chloride (3:1:0·005) and an aqueous wash; XMI, extracted with a mixture of xylene, methanol and isopropanol (1:2:4) without an aqueous wash.

These experiments suggested that DMT with an aqueous wash and XMI without an aqueous wash may offer similar overall facility for lipid extraction. We therefore undertook a deeper analysis of these two methods, using yet higher statistical power (*n* = 20 measurements) on three commercially-available animal milks and one preparation made from a nut (sold as almond drink). These measurements showed that XMI showed either the same or higher signal strength in all four sample types in both positive ionisation and negative ionisation modes, than DMT. The number of variables identified was also higher for XMI extracts in all animal milks. This trend was reversed for the almond preparation with around 30-60% more variables in both positive and negative ionisation modes.

These measurements were also used for calculating the coefficient of variance (CV). CVs were calculated based on variables that had 50% or more non-zero values. This range accommodates the different lipid profiles of the different milk sources used. CVs for XMI and DMT were particularly different. In positive mode, up to 80% of variables from XMI lipid extracts had a CV of less than 50%, with up to 51% of variables having a CV of less than 10% (Table 2). This is in contrast to DMT extracts in which 35-55% of variables had a CV of less than 50%. These results are reflected in negative ionisation mode, with around two thirds of variables in XMI extracts having a CV below 50%, whereas about 20% of variables from DMT extracts have a CV below 50% (Table 3). This consistency between measurements of CV in positive and negative ionisation modes suggests that performance of XMI for both glyceride isoforms (principally TGs, ionising in positive mode) and anionic/zwitterionic phospholipids is similar and superior to DMT.

**Table 2.** Coefficient of variance of lipid variables from extracts from dried milk spots, extracted wither using DMT or XMI methods, measured in positive ionisation mode. Only lipid variables with fewer than 50% zero values were used, due to the relatively wide range of the lipid profiles used. Dried milk spots were prepared from fresh, commercially available unhomogenised or standard whole milk, whole caprine milk or almond drink. *n* = 20 measurements per group.

|  | Fraction of variables identified in positive ionisation mode (%) |  |  |  |  |  |  |  |
| --- | --- | --- | --- | --- | --- | --- | --- | --- |
|  | DMT extracts |  |  |  | XMI extracts |  |  |  |
|  | Almond drink | Jersey milk | Caprine whole | Bovine whole | Almond drink | Jersey milk | Caprine whole | Bovine whole |
| 0-10% | 0 | 0 | 0 | 0 | 24 | 46 | 45 | 51 |
| 10-20% | 1 | 1 | 1 | 21 | 5 | 8 | 11 | 20 |
| 20-30% | 0 | 2 | 13 | 17 | 5 | 5 | 11 | 5 |
| 30-50% | 34 | 36 | 34 | 15 | 5 | 15 | 13 | 6 |
| >50% | 64 | 61 | 51 | 46 | 60 | 26 | 20 | 18 |
|  | 100%, 203 variables |  |  |  |  |  |  |  |

**Table 3.** Coefficient of variance of lipid variables from extracts from dried milk spots, extracted wither using DMT or XMI methods, measured in negative ionisation mode. Only lipid variables with fewer than 50% zero values were used. Dried milk spots were prepared from fresh, commercially available unhomogenised or standard whole milk, whole caprine milk or almond drink. *n* = 20 measurements per group.

|  | Fraction of variables identified in negative ionisation mode (%) |  |  |  |  |  |  |  |
| --- | --- | --- | --- | --- | --- | --- | --- | --- |
|  | DMT extracts |  |  |  | XMI extracts |  |  |  |
|  | Almond drink | Jersey milk | Caprine whole | Bovine whole | Almond drink | Jersey milk | Caprine whole | Bovine whole |
| 0-10% | 5 | 3 | 1 | 4 | 18 | 10 | 10 | 10 |
| 10-20% | 1 | 1 | 2 | 1 | 9 | 3 | 4 | 11 |
| 20-30% | 3 | 0 | 2 | 0 | 13 | 11 | 16 | 12 |
| 30-50% | 11 | 8 | 10 | 15 | 30 | 27 | 41 | 30 |
| >50% | 80 | 88 | 85 | 80 | 29 | 48 | 28 | 36 |
|  | 100%, 99 variables |  |  |  |  |  |  |  |

## Discussion

This study was motivated by the increasing interest in the use of DBS and DMS for studies of lipids in human growth and metabolism. Currently more lipid work has been done on DBSs, both in terms of the number of reports of studies using this sample type for lipidomics^1, 6, 7, 16^ (review^15^) and also reports of the analytical limits of this sample type^5, 17, 18, 30, 31^. DBSs and DMSs are attractive for clinical studies as they are a relatively non-invasive way to collect samples, and are easier to store and transport than fresh serum or plasma samples. Typically, liquid samples require more expensive consumables and labelling than DBS/DMS, and are transported and stored at −80 °C. However, the DBS as a means for storing material is not without its problems. It has been discovered that like fresh samples in storage, DBSs degrade with time. This can make results from older samples varied and unreliable. Degradation through oxidation and hydrolysis is particularly fast at room temperature^5^. However, it has also been found that degradation has a roughly linear relationship with respect to time and storage temperature. Such degradation is minimised where samples are stored at −80 °C, including over long periods (5y). In this study, we found that both the number of lipid variables and the total signal strength was higher in lipid extracts from DBSs treated with GCTU for 24h before extraction. We also found that a mixture of solvents with a spectrum of polarities was both efficient at extracting the lipid fraction from DMS and convenient for handling large numbers of small-scale extractions simultaneously (high throughput lipidomics).

An understanding of the precision of lipid extraction procedures is important in order to characterise and minimise batch effects between both individual DBS samples and studies. For example, it could be used to inform interpretation of increases in the abundance of *lyso*-lipids associated with particular phenotype(s). Cancer, metabolic disease and pregnancy all involve changes in the abundance of *lyso*-PC in the circulation. If these conditions were to be investigated using DBSs, false positive results could be avoided.

We found that both the number and strength of signals associated with lipid variables increased with the appropriate treatment of DBSs. This has also allowed us to assess the quality of different solvent systems used for extracting the lipid fraction from DMSs. Further checks could be made using orthogonal methods for determining molecular structure, for example through NMR. Dual spectroscopy^22^ has been designed as way of using orthogonal methods of spectroscopy, despite ^31^P NMR not being amenable to high throughput sample runs. Further work could include the use of ^1^H NMR for this purpose, however this has not been attempted to DBS or DMS-based samples yet, to our knowledge.

The 3D structure of DMSs suggested to us before commencing the study that they too may have a partially inaccessible lipid fraction. However, testing this hypothesis showed that the lipid fraction could be isolated well not only without powerful chaeotropes such as guanidine and thiourea, but even without water. This work therefore adds to formative work on extracting lipids from DMSs already reported^19^. Remarkably, the solvent system that proved to be best for extracting lipids from DMSs (xylene/methanol/isopropanol) contrasts sharply with the mixture of dichloromethane, methanol and triethylammonium chloride that is superior for extracting lipids from wet biological samples both in high throughput^21, 22, 25^ and large-sample-size ^32^ formats. This is based on calculations of the coefficient of variance that showed that the XMI method is generally more precise for extracting lipids from DMS. This is surprising as the number of variables and the total signal strength appear to be comparable between the DMT and XMI methods. Furthermore, the CVs for DMT extracts from DBSs are considerably better than those from DMSs. This set of observations suggests to us that dried spot sample matrices have distinct chemistries that benefit from tailored handling for lipid extraction.

This study was made possible by the realisation that high throughput mass spectrometry can be used to characterise both the number of lipids present and the amount (through signal strength). Previous discussion has noted that several studies have attempted to compare lipid extraction methods^23^. Typically, such studies have used samples with a large (>=2g lipid/sample) or medium (0·1-1·0g lipid/sample) scale sample set and profiled the extracts at lipid class level using chromatography. However, current requirements of lipidomics studies are for quantification of individual isoforms of lipids as well as class abundance, and for consistent precision in the measurement of all components. Such studies therefore suffer from a relatively weak measure of the quality of the extractions, coupled with a poor measure of variance (typically *n* = 3 extractions or fewer). In this study we have sought to address these concerns by using considerably more repeated measurements (*n* = 16-20 per group) and a more reliable form of structural identification that is established in identifying individual isoforms in a high throughput manner (direct infusion mass spectrometry). The striking contrast in precision between DMT and XMI methods, as shown by difference in CVs (Tables 2 and 3) suggest that calculation of CV is also a key part of assessing the efficiency of lipid extraction.

## Conclusion

This study was based on the hypotheses that exposure to chaeotropes would facilitate extraction of lipids from DBSs and DMSs, and that an extraction protocol without an aqueous wash was a more efficient method for isolating the lipid fraction from DMSs. It was motivated by the desire to improve the extraction efficiency for lipidomics studies. We conclude that exposure of DBSs to an aqueous solution of chaeotropes (GCTU) offers an improvement to lipidomics using this sample type, but that this is not required for DMSs. We also conclude that a non-aqueous method using a mixture of xylene, methanol and isopropanol can be used to extract lipids from DMSs efficiently, and that this can be used conveniently on high throughput lipidomics projects. These analyses lead us to conclude that dried spot matrices differ considerably in their handling requirements for extracting lipids.

## Acknowledgements

The authors would like to thank Drs S. G. Snowden, B. J. Jenkins, J. Warner and D. Parkington for helpful discussions and laboratory assistance. We gratefully acknowledge financial support from the BBSRC (BB/M027252/1) for SF.

## Supplementary Tables

**Supplementary Table 1.**
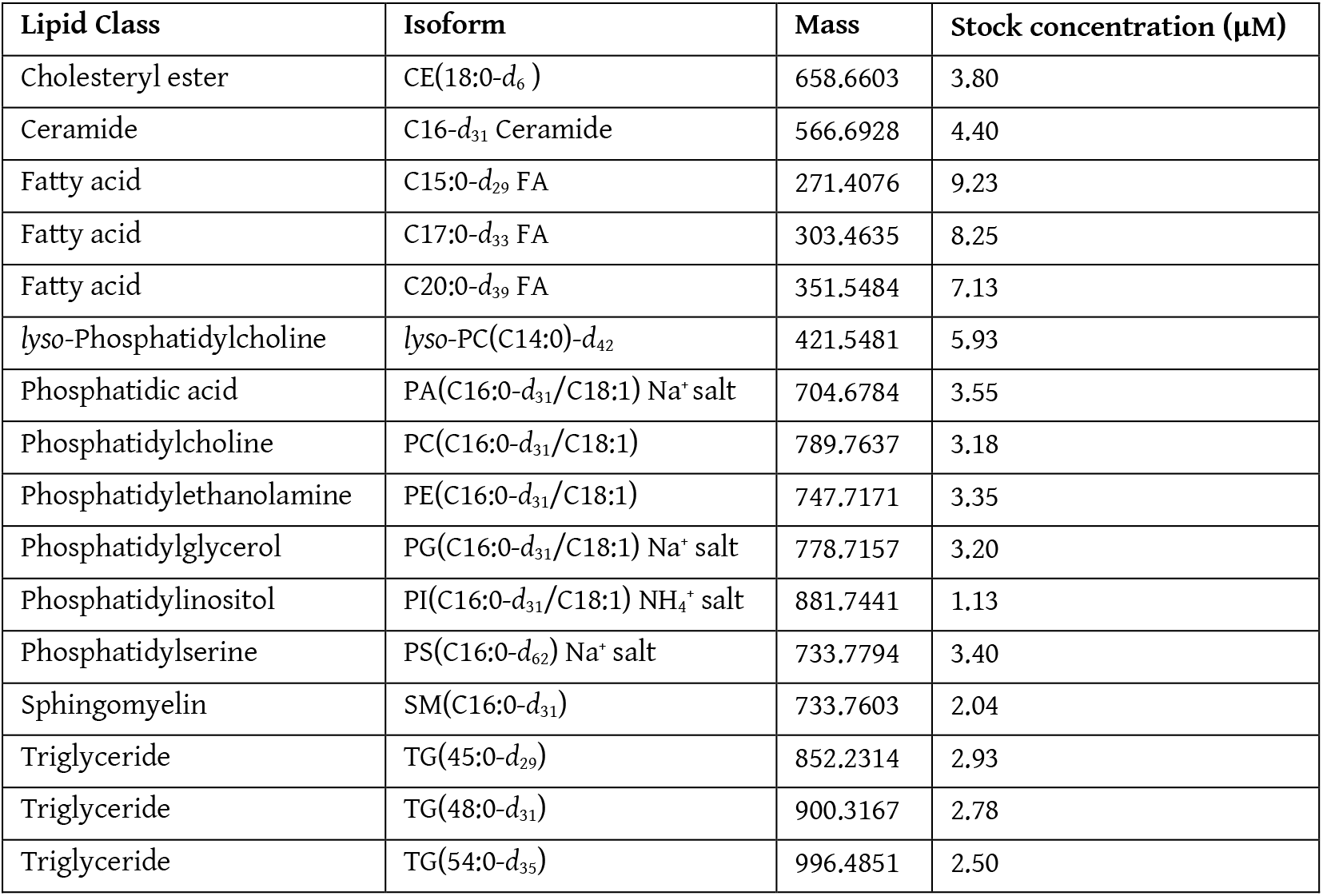
List of internal standards used for lipid profiling in the present study

| Lipid Class | Isoform | Mass | Stock concentration (μM) |
| --- | --- | --- | --- |
| Cholesteryl ester | CE(18:0-*d*<sub>6</sub>) | 658.6603 | 3.80 |
| Ceramide | C16-*d*<sub>31</sub> Ceramide | 566.6928 | 4.40 |
| Fatty acid | C15:0-*d*<sub>29</sub> FA | 271.4076 | 9.23 |
| Fatty acid | C17:0-*d*<sub>33</sub> FA | 303.4635 | 8.25 |
| Fatty acid | C20:0-*d*<sub>39</sub> FA | 351.5484 | 7.13 |
| lyso-Phosphatidylcholine | lyso-PC(C14:0)-*d*<sub>42</sub> | 421.5481 | 5.93 |
| Phosphatidic acid | PA(C16:0-*d*<sub>31</sub>/C18:1) Na<sup>+</sup> salt | 704.6784 | 3.55 |
| Phosphatidylcholine | PC(C16:0-*d*<sub>31</sub>/C18:1) | 789.7637 | 3.18 |
| Phosphatidylethanolamine | PE(C16:0-*d*<sub>31</sub>/C18:1) | 747.7171 | 3.35 |
| Phosphatidylglycerol | PG(C16:0-*d*<sub>31</sub>/C18:1) Na<sup>+</sup> salt | 778.7157 | 3.20 |
| Phosphatidylinositol | PI(C16:0-*d*<sub>31</sub>/C18:1) NH<sub>4</sub><sup>+</sup> salt | 881.7441 | 1.13 |
| Phosphatidylserine | PS(C16:0-*d*<sub>62</sub>) Na<sup>+</sup> salt | 733.7794 | 3.40 |
| Sphingomyelin | SM(C16:0-*d*<sub>31</sub>) | 733.7603 | 2.04 |
| Triglyceride | TG(45:0-*d*<sub>29</sub>) | 852.2314 | 2.93 |
| Triglyceride | TG(48:0-*d*<sub>31</sub>) | 900.3167 | 2.78 |
| Triglyceride | TG(54:0-*d*<sub>35</sub>) | 996.4851 | 2.50 |

